# Integrative Transcription Start Site Analysis and Physiological Phenotyping Reveal Torpor-specific Expressions in Mouse Skeletal Muscle

**DOI:** 10.1101/374975

**Authors:** Genshiro A Sunagawa, Ruslan Deviatiiarov, Kiyomi Ishikawa, Guzel Gazizova, Oleg Gusev, Masayo Takahashi

## Abstract

Mice enter an active hypometabolic state, called daily torpor, when they experience a lowered caloric intake under cool ambient temperature (*T*_*A*_). During torpor, the oxygen consumption rate (*VO*_*2*_) drops to less than 30% of the normal rate without harming the body. This safe but severe reduction in metabolism is attractive for various clinical applications; however, the mechanism and molecules involved are unclear. Therefore, here we systematically analyzed the expression landscape of transcription start sites (TSS) in mouse skeletal muscles under various metabolic states to identify torpor-specific transcription patterns. We analyzed the soleus muscles from 38 mice in torpid, non-torpid, and torpor-deprived conditions, and identified 287 torpor-specific promoters. Furthermore, we found that the transcription factor ATF3 was highly expressed during torpor deprivation and that the ATF3-binding motif was enriched in torpor-specific promoters. Our results demonstrate that the mouse torpor has a distinct hereditary genetic background and its peripheral tissues are useful for studying active hypometabolism.

## INTRODUCTION

Mammals in hibernation or in daily torpor reduce their metabolic rate to 1-30% of that of euthermic states and enter a hypothermic condition without any obvious signs of tissue injury (Bouma et al., 2012; Geiser, 2004). How mammals adapt to such a low metabolic rate and low body temperature without damage remains as one of the central questions in biology. Mammals maintain their *T*_*B*_ within a certain range by producing heat. In cold, the oxygen requirements for heat production increases, at a rate negatively proportional to the body size (Heldmaier et al., 2004). Instead of paying the high cost for heat production, some mammals are able to lower their metabolism by sacrificing body temperature homeostasis. This condition, in which the animal reduces its metabolic rate followed by whole-body hypothermia, is called active hypometabolism. As a result, the homeostatic regulation of body temperature is suppressed, and the total energy usage is spared. This hypometabolic condition is called hibernation when it lasts for a season, and daily torpor when it occurs daily.

Four conditions have been proposed to occur in active hypometabolism in mammals (Sunagawa and Takahashi, 2016): 1) resistance to low temperature, 2) resistance to low oxygen supply, 3) suppression of body temperature homeostasis, and 4) heat production ability under a low metabolic rate. Of these conditions, 1) and 2) were found to be cell/tissue-specific or local functions, which prompted researchers to analyze genome-wide molecular changes in various tissues of hibernators, including brain, liver, heart, skeletal muscles, and adipose tissues. A major role of differential gene expression in the molecular regulation of hibernation was first suggested by Srere with co-authors, who demonstrated both mRNA and protein upregulation of α2-macroglobulin during torpor in the plasma and liver of two ground squirrel species (Srere et al., 1995). With the development of high-throughput sequencing approaches, such as RNA-seq and microarrays, series of transcriptomic investigations were conducted in well-studied hibernating animals, including ground squirrels (Hampton et al., 2011; Schwartz et al., 2013; Williams et al., 2005), bears (Fedorov et al., 2009, 2014; Zhao et al., 2010), and bats (Lei et al., 2014; Seim et al., 2013). Recent proteomics studies in ground squirrels using two-dimensional gel electrophoresis (Epperson et al., 2004; Martin et al., 2008) and shotgun proteomics (Shao et al., 2010) also explored the post-transcriptional regulation of hibernation. Furthermore, several studies demonstrated a strict epigenetic control of hibernation (Alvarado et al., 2015; Biggar and Storey, 2014), and a role of miRNAs in the process (Chen et al., 2013; 5 Luu et al., 2016).

At the same time, due to the lack of detailed genome information abouthibernators, i.e., squirrels, bats, and bears, the interpretation of high-throughput sequencingresults is challenging. Instead, the mouse, *Mus musculus*, has rich genetic resources andcould overcome this weakness. One noteworthy feature of this animal is the abundantvariety of inbred strains and the diverse phenotypes. For example, the sleep phenotype (Franken et al., 1998; Koehl et al., 2003), circadian phenotype (Kopp, 2001; Schwartz andZimmerman, 1990), cocaine response (Ruth et al., 1988), and reproductive system (Mochida et al., 2014) are well-known to show differences among inbred strains. Moreover,recent advances in genetic engineering make mice even more attractive for genetictweaking at the organismal level. Transgenic animals in the F_0_ generation are used to testgenetic perturbation directly at the organismal level phenotypes in mice (Sunagawa et al.,2016; Wang et al., 2013). Notably, the mouse is well-known to enter daily torpor (Hudsonand Scott, 1979), and we recently developed a method to reproducibly inducing torpor inmice (Sunagawa and Takahashi, 2016), making the mouse a suitable animal for studyinghypometabolism.

The goal of this study was to analyze contribution of the genetic background to thetorpor phenotype by introducing the mouse as model for active hypometabolism, takingadvantage of the rich and powerful genetic technologies available for this animal. First, wefound that two genetically close mice inbred strains, C57BL/6J (B6J) and C57BL/6NJcl(B6N), exhibit distinct torpor phenotypes; B6N has a higher metabolism during torpor and alower rate of torpor entry. To clarify the genetic link to the mouse torpor phenotype, weperformed Cap Analysis of Gene Expression (CAGE) in soleus muscles taken from 38animals under various metabolic conditions. We found that entering torpor and restoringactivity were associated with distinct changes in the transcriptomic profile, including markedchanges in promoter shapes. Finally, we present evidence that the torpor-specificpromoters are related to the genetic differences in these inbred strains.

## RESULTS

### Torpor Phenotype in Mice is Affected by Genetic Background

We previously showed that 100% of B6J mice enter daily torpor when they are deprived of food for 24 hours under a *T*_*A*_ of 12 to 24 °C (Sunagawa and Takahashi, 2016). B6N is an inbred strain that is genetically close to B6J (Simon et al., 2013). Despite their genetic resemblance, these two strains show significant differences in their sleep time (Sunagawa et al., 2016), reaction to cocaine (Kumar et al., 2013), and energy expenditure (Simon et al.,2013). Therefore, we decided to compare the daily torpor phenotype between these two strains.

First, we tested whether the torpor induction method developed for B6J could be applied to B6N. We set a B6N mouse into the test chamber on day 0 and used the 72-hour-data from the beginning of day 1 for analysis. Keeping the animal in a constant *T*_*A*_, weremoved the food for 24 hours from the beginning of day 2 (Figure 1A). *T*_*B*_ and *VO*_*2*_ weresimultaneously recorded for 72 hours, and the first 24 hours of data were used to estimate the individual basal metabolism of the animal. The metabolism was evaluated every 6minutes, and when it was lower than the estimated baseline, the animal was defined to be in a low-metabolism condition. In this study, when the animal showed low metabolism during the latter half of the second day, which is the dark phase in which mice are normally active, the state was labeled as “torpor”. Figure 1B shows a representative torpor pattern ofa male B6N mouse. We tested the torpor entry rate of B6N mice by inducing torpor atvarious *T*_*A*_s, (8, 12, 16, 20, 24 and 28 °C) (Figures S1A and S1B). B6N mice entered torpor at a peak rate of 100% at *T*_*A*_ = 16 °C, but the rate decreased at higher or lower *T*_*A*_s (Figure1C). Notably, at *T*_*A*_ = 8 °C, more than 60% of the animals died without entering torpor. The overall entry rates were lower than those of B6J, which enter torpor at 100% when *T*_*A*_ = 12to 24 °C (Sunagawa and Takahashi, 2016).

**Figure 1.**
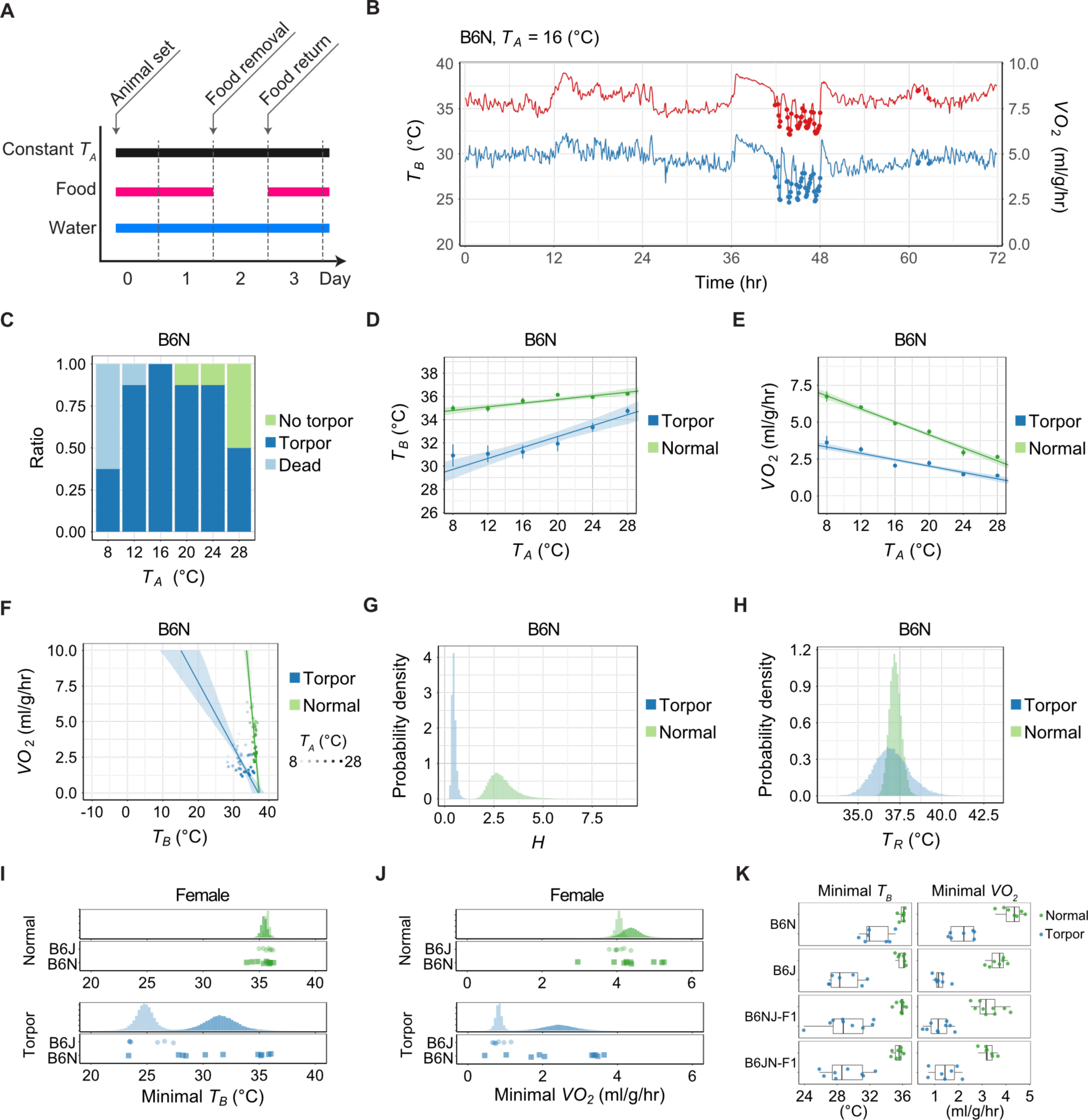
Torpor Phenotype is Affected by Genetic Background. (A) Protocol for fasting-induced daily torpor in a mouse. (B) Representative metabolic transition of mouse daily torpor. Red and blue lines denote *T*_*B*_and *VO*_*2*_, respectively. Filled circles on the line are time points evaluated as “torpor”. (C) Torpor entry rate in the male B6N mouse. It peaked at *T*_*A*_ = 16 °C. (D) (E) Minimal *T*_*B*_ and *VO*_*2*_ of male B6N at various *T*_*A*_s. In these and the following panels,green and blue denote the normal and torpid states, respectively. Dots with the verticalerror bars denote the observed mean and SEM of the minimal variables [*T*_*B*_ in (D), *VO*_*2*_ in(E)] at each *T*_*A*_, and the line and shaded area denote the mean and the 89% HPDI intervalsof the estimated minimal variables. (F) Relationship between minimal *T*_*B*_ and *VO*_*2*_ seen during normal and torpid states atvarious *T*_*A*_s. Darkness of the dot indicates the *T*_*A*_. The horizontal intercept of the lineindicates the theoretical set-point of *T*_*B*_, which is *T*_*R*_. In the normal state, *T*_*B*_ is kept relativelyconstant by using oxygen and producing heat to fill the gap between *T*_*R*_ and *T*_*A*_. On theother hand, during daily torpor, the sensitivity against *T*_*B*_ is weakened, which is visualizedby a less steep slope. (G) Posterior distribution of the estimated *H* during the normal state and torpor. The lack ofoverlap strongly suggests that *H* is different between these two conditions. (H) Posterior distribution of the estimated *T*_*R*_ during normal state and torpor. The highoverlap of the two distributions suggests that *T*_*R*_ is indistinguishable between these twoconditions. (I) (J) Minimal *T*_*B*_ and *VO*_*2*_ of female B6N at *T*_*A*_ = 20 °C. Lighter color is B6J, and darker isB6N. Upper panel shows the posterior distribution of the estimated minimal *T*_*B*_ and *VO*_*2*_,and the lower panel shows the raw data for each group. (K) Minimal *T*_*B*_ and *VO*_*2*_ of male mice at *T*_*A*_ = 20 °C. B6NJ-F1 is the offspring of a B6Nmother and B6J father. Note that the higher metabolic rate in the torpor phenotype of B6Ndisappears when crossed with B6J.

We next examined how the metabolism varied with the *T*_*A*_ changes in B6N mice.Specifically, the minimal *VO*_*2*_ and the minimal *T*_*B*_ during normal and torpid states overvarious *T*_*A*_s were entered into a statistical model to obtain the posterior distributions of parameters of the thermoregulatory system (Sunagawa and Takahashi, 2016). Because mice are homeothermic, the minimal *T*_*B*_ under normal conditions was expected to beunaffected by *T*_*A*_. We calculated the unit-less slope *a*_*1*_ defined by the change in *T*_*B*_ againstunit *T*_*A*_ change. When a specific total probability α is given, the highest posterior densityinterval (HDPI) is defined by the interval of a probability density, which includes values morecredible than outside the interval and the total probability of the interval being α. In thiscase, the given dataset was predicted to have an 89% HDPI of *a*_*1*_ as [0.054, 0.103](Figures 1D and S1C; hereafter, 89% HDPI will be indicated by two numbers in squarebrackets.). Thus, under normal conditions, *T*_*B*_ had a very low sensitivity against *T*_*A*_; when *T*_*A*_changed 10 degrees, the *T*_*B*_ changed no more than 0.5 to 1 degree. In contrast, duringtorpor, the minimal *T*_*B*_ was more sensitive to *T*_*A*_, which was described by a larger *a*_*1*_ thanunder normal conditions (*a*_*1*_ during torpor was [0.166, 0.312]; Figures 1D and S1C). The*VO*_*2*_ also differed between the normal and torpid conditions in B6N. In a homeothermicanimal, *VO*_*2*_ decreases when *T*_*A*_ increases because less energy is needed for heatproduction. This was, indeed the case in B6N mice (Figure 1E). The slope *a*_*2*_ (ml/g/hr/°C),defined by the negative change in *VO*_*2*_ against unit *T*_*A*_ change, was estimated to be [0.199,0.239] ml/g/hr/°C under normal conditions (Figure S1D). During torpor, however, theanimals reduced their *VO*_*2*_ to nearly half of value under the normal condition (*a*_*2*_ duringtorpor was [0.091, 0.126] ml/g/hr/°C; Figures 1E and S1D). Thus, both *T*_*B*_ and *VO*_*2*_ showedhypometabolic transitions during torpor in B6N mice.

To compare the function of the heat-production system between B6J and B6N, weestimated the negative feedback gain (*H*) and the theoretical target temperature (*T*_*R*_) (°C) ofB6N from the *VO*_*2*_ and *T*_*B*_ observed at various *T*_*A*_s. As it was described in our previousstudy (Sunagawa and Takahashi, 2016), we applied the recorded *VO*_*2*_ and *T*_*B*_ to a statisticalmodel. The estimated median *H* dropped 83.8% during torpor (Figures 1F and 1G) whilethe *T*_*R*_ dropped slightly (the estimated median *T*_*R*_ difference from normal to torpor was0.25 °C; Figures 1F and 1H). To compare these parameters with B6J mice, we recorded the*T*_*A*_ = 28 °C data missing in our previous study and recalculated both the *H* and *T*_*R*_ for B6Jmice using *T*_*A*_s of 8, 12, 16, 20, 24 and 28 °C (Figure S1E). In this case, they showed a94.0% drop in the estimated median *H* and 0.68 °C drop in *T*_*R*_ during torpor. Based on the estimated distributions, during torpor, B6N mice had a smaller *H* than B6J (*ΔH* was [0.020,0.366], which was totally positive; Figure S1F). Interestingly, Δ*T*_*R*_ during torpor was [-1.47,3.33] °C, which included zero in the 89% HDPI, indicating that the difference between B6Jand B6N was not clear in these groups (Figure S1G).

To confirm that the phenotype difference between B6N and B6J was not sex-specific, we recorded the torpor phenotypes of female B6J and female B6N at *T*_*A*_ = 20 °C (Figures S1H and S1I). As observed in males, the females showed similar minimal *T*_*R*_ and*VO*_*2*_ under the normal condition, and B6N showed a higher metabolic rate during torporthan B6J. During torpor, the estimated minimal *T*_*B*_ was [29.1, 33.9] °C and [23.4, 26.1] °Cand minimal *VO*_*2*_ was [1.72, 3.12] ml/g/hr and [0.68, 0.96] ml/g/hr in B6N and B6J mice,respectively (Figures 1I and 1J). The posterior distribution of differences in the minimal *T*_*B*_and *VO*_*2*_ during torpor from B6J to B6N was [3.96, 9.56] °C and [0.89, 2.31] ml/g/hr,respectively. Both 89% HPDIs were greater than zero, which mean the probability that B6Nhas a higher minimal *T*_*B*_ and *VO*_*2*_ during torpor than B6J is greater than 89%.

Because inbred strains have an identical genomic background, our results stronglyindicate that the torpor phenotype is related to the genomic difference between these twoinbred strains. To examine this possibility, we crossed B6J and B6N and evaluated thetorpor phenotype of their offspring. We performed two types of mating combinations: femaleB6N with male B6J (B6NJ-F1) and female B6J with male B6N (B6JN-F1). In bothcombinations, the F1 generations showed the B6J phenotype during torpor (Figure 1K). Both the minimal *T*_*B*_ and the minimal *VO*_*2*_ during torpor of B6J, B6NJ-F1, and B6JN-F1were lower than that of B6N (Figure S1J).

All of these results indicated that the hypometabolic phenotype during torpor isinheritable. Interestingly, B6N and B6J only have 140,111 SNP bases (Keane et al., 2011). We next examined the hypothesis, that there should be certain genetic variation in twostrains associated with genes or regulatory elements contributing to control of torpor. Because B6N and B6J did not have a difference in *T*_*R*_ (Figures 1F and S1E), which isregulated in the thermoregulatory center located in the hypothalamus (Nakamura, 2011), we hypothesized that the phenotypic difference between B6N and B6J occurs in the peripheral tissue. Therefore, we used peripheral muscle to test for torpor-specific RNA expressions and to identify the responsible genetic network for torpor.

### Fasting-induced Torpor Shows a Reversible Transcriptome Signature

Animals in active hypometabolism return to the normal condition without any damage even after experiencing extreme hypothermic and hypometabolic conditions. To analyze reversibility in peripheral tissue gene expression during torpor, we isolated soleus muscles on day 1 (Pre, n = 4), 2 (Mid, n = 8) and 3 (Post, n = 4) at ZT-22 as experiment #1 (Figure2A). We chose these time points because B6J mice usually start to enter torpor at around ZT-14, and at ZT-22, which is two hours before the light is turned on, the animals are very likely to be in a torpid state (Sunagawa and Takahashi, 2016). Indeed, the *VO*_*2*_ was higherin the Pre and Post groups, and was lowest in the Mid group (Figure 2B). Skeletal muscle isa popular tissue for hibernation research because they show little atrophy even during prolonged immobility. Therefore, a considerable number of transcriptomic and proteomic studies have been performed in the past (Bogren et al., 2017; Fedorov et al., 2014;Hampton et al., 2011; Hindle and Karimpour-Fard, 2011; Muleme et al., 2006), which encouraged us to choose skeletal muscle as a target tissue. We extracted the RNA from the muscle samples, and performed single-molecule sequencing combined with CAGE (Kanamori-Katayama et al., 2011; Kodzius et al., 2006). This method allowed us to evaluate the genome-wide distribution and quantification of TSSs in these tissues. Based on the TSS distribution, we identified 12,862 total CAGE clusters (promoters). Among all the promoters,11,133 were associated with 10,615 genes.

**Figure 2.**
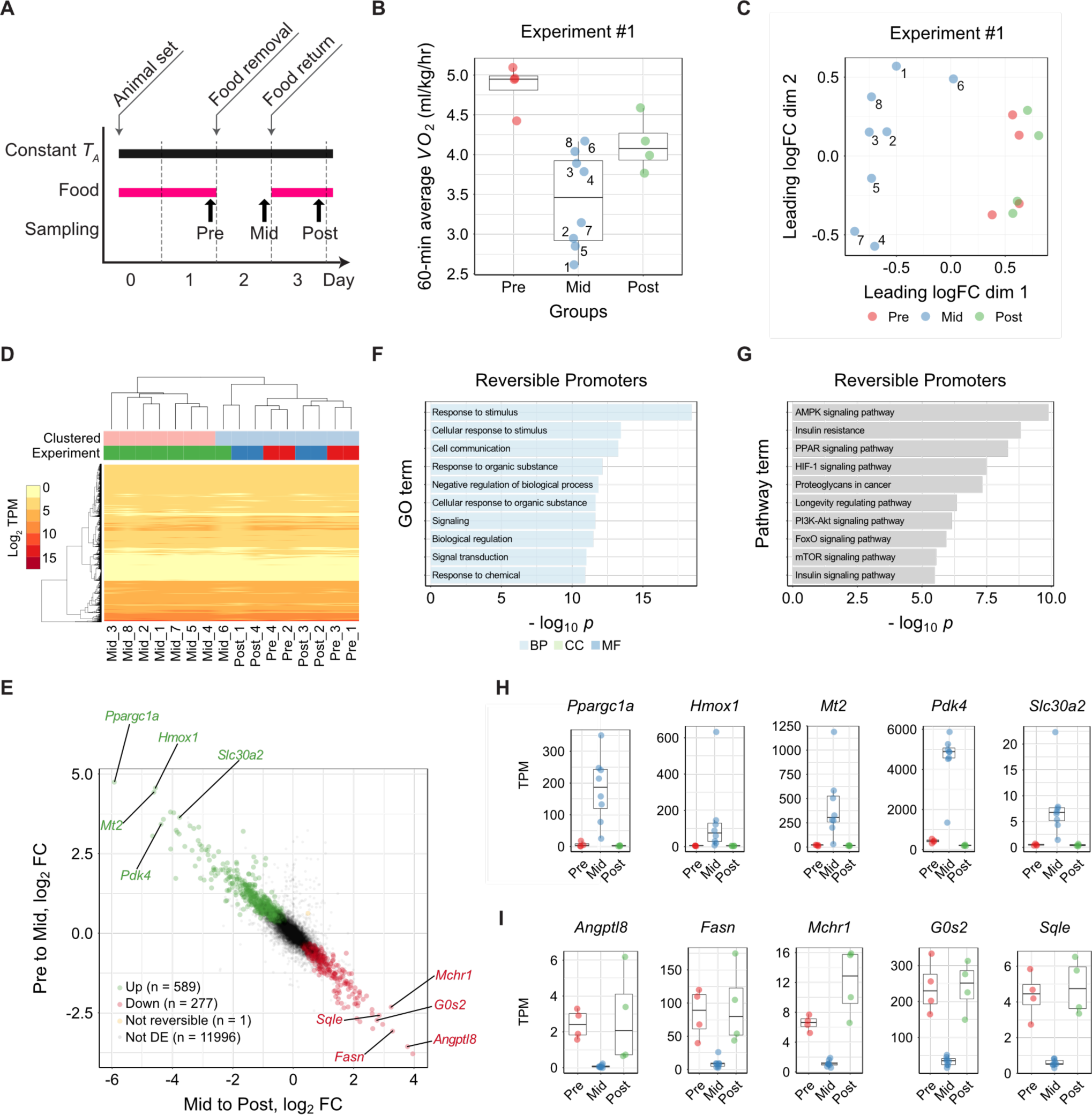
Fasting-induced Torpor Shows a Reversible Transcriptome Signature. (A) Protocol for sampling muscles from Pre, Mid, and Post torpor animals to test thereversibility of the transcriptional profile of muscles during torpor. (B) Boxplots for the *VO*_*2*_ of animals at sampling in the reversibility experiment #1. Each dotrepresents one sample from one animal. During torpor (Mid group), the median *VO*_*2*_ waslower than during Pre or Post torpor. The band inside the box, the bottom of the box, and the top of the box represent the median, the first quartile (Q_1_), and the third quartile (Q_3_),respectively. The interquartile range (IQR) is defined as the difference between Q_1_ and Q_3_. The end of the lower whisker is the lowest value still within 1.5 IQR of Q_1_, and the end of the upper whisker is the highest value still within 1.5 IQR of Q_3_. Every other boxplot in this manuscript follows the same annotation rules. The numbers in the Mid torpor group are identification numbers of the animals. (C) MDS plot of the TSS-based distance in reversibility experiment #1. Each dot represents one sample from one animal. The Mid group clustered differently from the Pre and Post groups in the 1st dimension. The two internal groups seen in the Mid group in Figure 2B were not evident in this plot, indicating the transient metabolic change during torpor was not correlated with transcription. (D) Hierarchical clustering heatmap based on the TPM of TSS detected in the reversibility experiment #1. (E) Distribution of CAGE clusters according to the fold-change in the TPM of Pre to Mid and Mid to Post torpor. The top five up- and down-regulated reversible promoters that had annotated downstream genes are shown. (F) Top ten enriched GO terms in the reversible promoters. (G) Top ten enriched KEGG pathways in the reversible promoters. (H) (I) Top five up- and down-regulated reversible promoters ordered according to the magnitude of the TPM change. Promoters that had annotated downstream genes areshown.

The multidimensional scaling (MDS) plot of the promoter-level RNA expression showed that the Pre and Post groups had distinct expression profiles from the Mid group (Figures 2C and 2D). During torpor, the animal may show both high and low metabolism due to the oscillatory nature of this condition (note the wavy pattern of *VO*_*2*_ in Figure 1B).Indeed, the animal in the Mid group showed a broad diversity of metabolic rates (Figure 2B). Each number in Figures 2B and 2C represents the same animal in the Mid group. Despite the broad metabolism range during torpor (Figure 2B), the CAGE cluster profile did not show clustering within the Mid group according to metabolic state (Figures 2C and 2D), indicating that the hourly oscillatory change in metabolism during torpor is based on atranscription-independent mechanism.

To test the reproducibility of this experiment, we performed another independent set of samplings and CAGE analysis (experiment #2). We obtained 2, 5, and 3 samples forthe Pre, Mid, and Post states, respectively. In experiment #2, the *VO*_*2*_ at sampling showeda similar pattern as in experiment #1 (Figure S2A), and the MDS plot showed that the Preand Post groups had a distinct transcriptome profile from the Mid group (Figures S2B and S2C). These results were consistent with those of experiment #1.

To gain insight into the biological process underlying the reversible expression during torpor, we analyzed differentially expressed (DE) genes on the level of promoters inthe Pre to Mid and in the Mid to Post conditions. The promoters were considered differentially expressed when the false discovery rate (FDR) was smaller than 0.05. Reversibly up-regulated DE promoters were defined if they show a significant increase from the Pre to Mid (FDR < 0.05) and decrease from the Mid to Post (FDR < 0.05). Reversibly down-regulated DE promoters were similarly defined but in the opposite direction (Figure2E). We found 589 up-regulated and 277 down-regulated promoters (representing 481 and221 genes) from the 12,863 total promoters, with enrichment in several distinct KEGG pathways. The top 10 enriched GO terms and KEGG pathways related to both the reversibly up- and down-regulated DE genes are shown in Figures 2F and 2G. Furthermore, we found enrichment of certain motifs in the promoters with reversible dynamics of expression (Figures S2D and S2E). Finally, every up- and down-regulated DE promoter was ranked in the order of the total fold-change, which was the sum of the fold-changes in both the Pre to Mid and the Mid to Post (Figures 2H and 2I).

To exclude the possibility that the difference we observed was the direct effect of starvation and not the low metabolism, we further analyzed the transcriptomic profile of mouse muscles under several conditions that can prevent the animal from entering torpor.

#### Torpor Prevention at High *T*_*A*_ Revealed Hypometabolism-associated Promoters

Torpor can be induced by removing food for 24 hours only when the animal is placed in arelatively low *T*_*A*_. We have shown that B6J mice enter torpor at a rate of 100% from *T*_*A*_ =12 °C up to *T*_*A*_ = 24 °C (Sunagawa and Takahashi, 2016) and that some animals stop entering torpor at *T*_*A*_ = 28 °C (Figures S3A and S1E). We further tested whether the animals could enter torpor at *T*_*A*_ = 32 °C (Figure S3B). In this warm condition, even if the animals were starved they did not enter torpor, possibly due to the lack of heat loss than at lower*T*_*A*_s. Taking these two requirements into account, fasting and low *T*_*A*_, we designed twotorpor-preventive conditions and compared the expression in the muscles under these conditions to that under the ideal torpor state (Figure 3A). One is a high *T*_*A*_ (HiT)environment and the other is a non-fasted (Fed) condition. Both conditions prevented the animals from inducing torpor, because the two essential requirements were lacking. We then, compared the tissue from these conditions to the ideal torpid tissue, which was from fasting animals at a low *T*_*A*_, and obtained the transcripts that were differentially expressed from torpor in each non-torpor condition. The expression differences shared in these two experiments would be those affected by both low *T*_*A*_ and fasting, and therefore would be the essential expressions for active hypometabolism, hereafter defined as hypometabolicpromoters.

**Figure 3.**
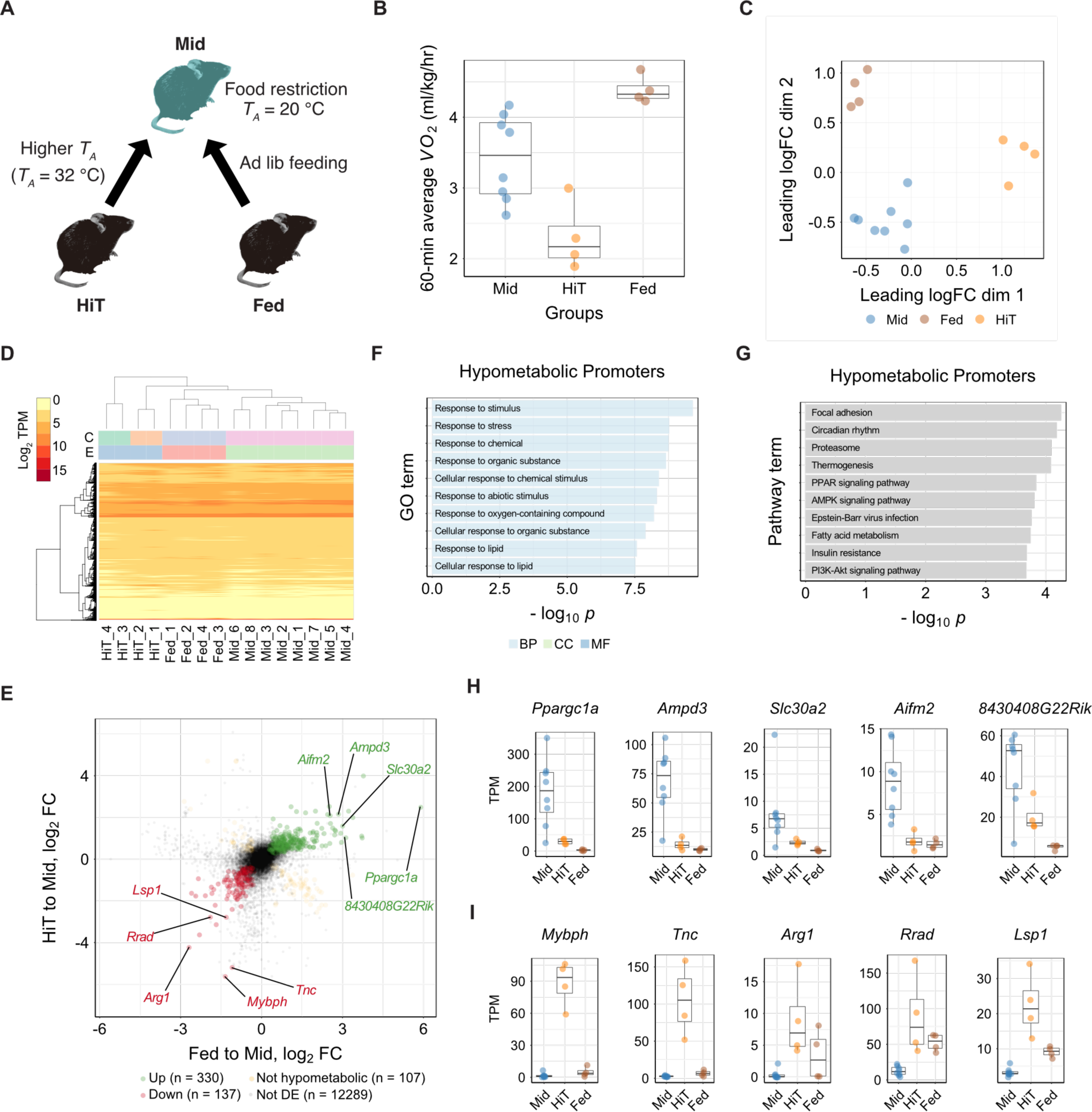
Torpor Prevention at high *T*_*A*_ Revealed Hypometabolism-associatedPromoters. (A) Protocol for detecting the hypometabolic expression by sampling muscles from twogroups in which torpor was prevented (HiT and Fed groups, n = 4 for each). For the torpid group, the samples collected in the reversibility test was used (Mid group, n = 8). (B) Boxplots for the *VO*_*2*_ of animals at sampling in the hypometabolic experiment. Each dot represents one sample from one animal. During torpor prevention by high-*T*_*A*_ (HiT group),the *VO*_*2*_ was lower than in the Mid group, and when torpor was prevented by food administration (Fed group), *VO*_*2*_ was higher than in the Mid group. (C) MDS plot of the TSS-based distance in the hypometabolic experiment. Each dot represents one sample from one animal. The Mid, Fed, and HiT groups were clustered separately. (D) Hierarchical clustering heatmap based on TPM of the TSS detected in the hypometabolic experiment. (E) Distribution of CAGE clusters according to the fold-change in TPM of the HiT to Mid and Fed to Mid groups. The top five up- and down-regulated hypometabolic promoters that hadannotated downstream genes are shown. (F) The top ten enriched GO terms in the hypometabolic promoters. (G) The top ten enriched KEGG pathways in the hypometabolic promoters. (H) (I) The top five up- and down-regulated hypometabolic promoters ordered according tothe magnitude of the TPM change. Promoters that had annotated downstream genes areshown.

We first compared the *VO*_*2*_ in the HiT and Fed groups against the Mid group (Figure 3B). Even though both groups had no animals entering torpor, the HiT group showed a lower *VO*_*2*_ while the Fed group showed a higher metabolism. Next, we compared the expression profile acquired from the CAGE analysis of tissues from both groups. TheMDS plot and hierarchical clustering showed that the Mid, Fed, and HiT groups consisted of independent clusters (Figures 3C and 3D). This finding indicated that the expressions during torpor (Mid group) were distinct from those during starvation alone (HiT) or at low *T*_*A*_ alone (Fed).

To extract the hypometabolic promoters, we performed the DE analysis (Figure 3E) between the HiT to Mid and the Fed to Mid. CAGE clusters up-regulated in both the HiT to Mid and the Fed to Mid were those that were upregulated during torpor regardless of theinitial condition, i.e., warm *T*_*A*_ or no fasting (green dots in Figure 3E). There were 330 of these up-regulated hypometabolic promoters from the total 12,863. On the other hand,CAGE clusters that were down-regulated in both the HiT to Mid and the Fed to Mid, werepromoters that were down-regulated regardless of the initial condition, and thus were thedown-regulated hypometabolic promoters (red dots in Figure 3E). The enrichment analysesof GO terms and KEGG pathways were performed (Figures 3F and 3G), and the motifsenriched in the hypometabolic promoters were also analyzed (Figures S3C and S3D). Thetop five promoters that had annotated genes nearby are listed as up- and down-regulatedhypometabolic promoters in Figures 3H and 3I, respectively.

These results showed that considerable numbers of genes are involved in theactive hypometabolic process independent from the responses to both hunger and cold. One of these genes, *Ppargc1a*, which was found at the top of the up-regulatedhypometabolic promoters, was also found at the top of up-regulated reversible promoters (Figure 2H). This is a good candidate for a torpor-specific gene, because it belongs to boththe reversible and the hypometabolic group in this study. Therefore, we next merged theresults of the reversible and the hypometabolic promoters to specify the torpor-specificpromoters and elucidate the fundamental transcriptional network of active hypometabolismin peripheral tissues.

### Identification of Torpor-specific Promoters and their Dynamics

Our two independent analyses, which focused on two essential torpor characteristics, i.e.,reversibility and hypometabolism, revealed that the skeletal muscle of torpid mice has aspecific transcriptomic pattern. Combining these results, we obtained torpor-specificpromoters, defined as the intersection of the reversible and the hypometabolic promoters. We found 226 up-regulated and 61 down-regulated torpor-specific promoters (Figure 4A). The top five promoters ordered according to the sum of the fold-change observed in the twogroups (reversible and hypometabolic promoters) are shown in Figures 4B and 4C. Remarkably, “protein binding” in the molecular function category in the GO terms was listedin the top ten enriched GO terms (Figure S4A). This group includes various protein-binding genes products, including transcription factors. To highlight the predominant transcriptional pathway related to torpor, we ran an enrichment study of KEGG pathways with the torpor-specific promoters. We obtained 13 pathways that showed statistically significant enrichment (Figure 4D). In particular, the mTOR pathway, which includes various metabolic processes related to both hibernation and starvation, was identified (Figure S4B). Furthermore, we analyzed the enriched motifs in the torpor-specific promoters (Figures S4C and S4D) and found 131 significantly enriched motifs out of 579 motifs registered inJASPAR 2018 (Khan et al., 2018).

**Figure 4.**
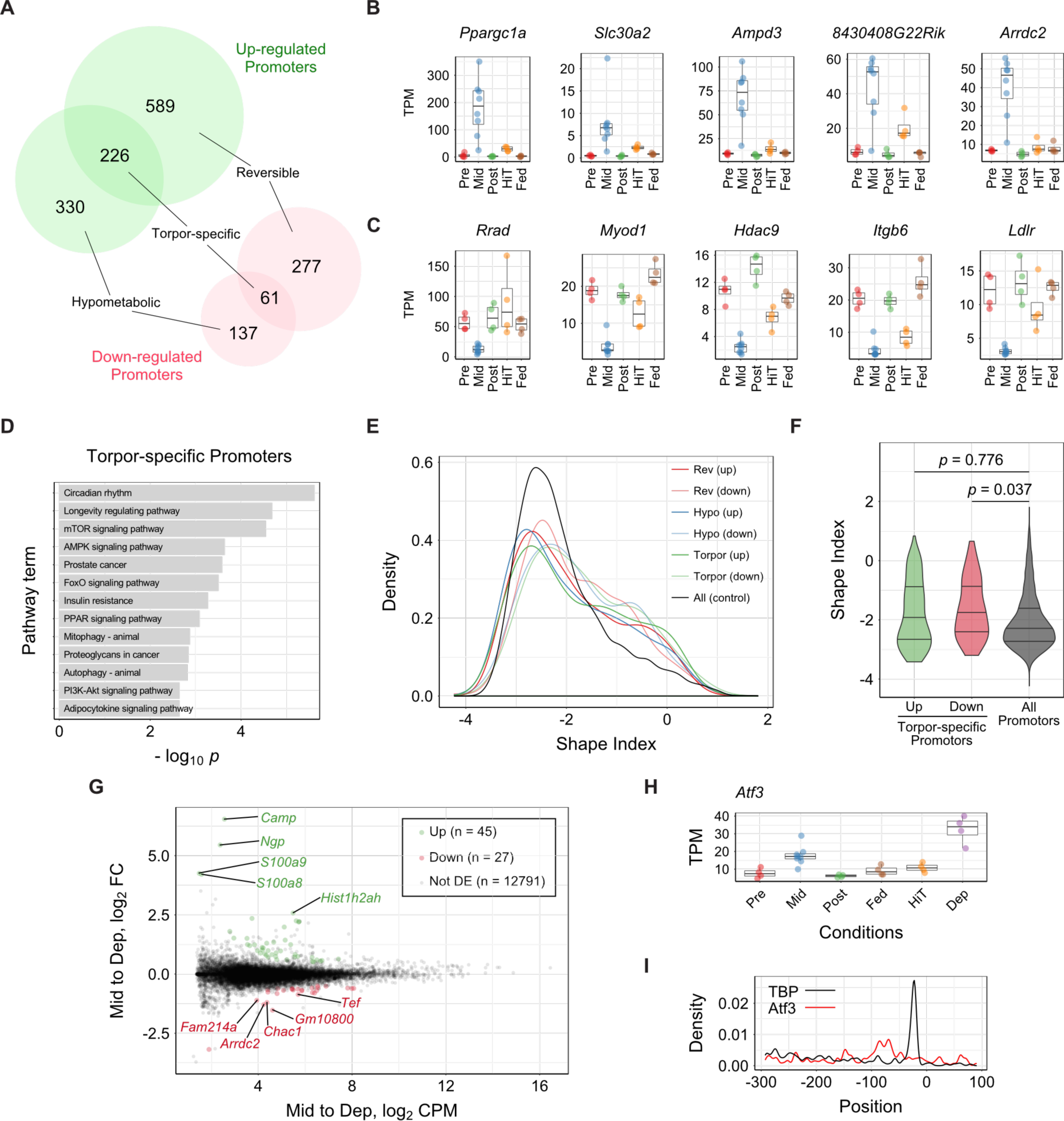
Identification of Torpor-specific Promoters and their Dynamics. (A) Torpor-specific promoters were defined by the intersection of reversible and hypometabolic promoters. Up-regulated torpor-specific promoters (n = 226), which were CAGE clusters that were highly expressed exclusively during torpor, were at the intersection of the up-regulated reversible (n = 589) and hypometabolic promoters (n =330). Down-regulated torpor-specific promoters (n = 61), which were CAGE clusters thatwere highly suppressed exclusively during torpor, were at the intersection of down-regulated reversible (n = 277) and hypometabolic promoters (n = 137). (B) (C) Top five up-regulated (B) and down-regulated (C) torpor-specific promoters orderedaccording to the sum of the TPM change observed in the reversibility and hypometabolismexperiments. Only promoters that had annotated downstream genes are shown. (D) Top ten enriched KEGG pathways in the torpor-specific promoters. (E) Distribution of the SI of all of the mouse muscle promoters. An SI of 2 indicates asingleton-shaped CAGE TSS signal, and promoters with SI < −1 have a broad shape. (F) Distribution of the SI for torpor-specific promoters compared to all muscle promoters. The three horizontal lines inside the violin denote the 1st, 2nd, and 3rd quartile of thedistribution from the upmost line. (G) Distribution of CAGE clusters according to the mean TPM and the fold-change TPM of the Mid to Dep group. Top five up- and down-regulated torpor-deprivation-specificpromoters that had annotated downstream genes are shown. (H) Among the torpor-specific up-regulated genes, *Atf3* was the only DE gene during torpordeprivation. (I) Enrichment profile of the ATF3-binding motif in torpor-specific promoter regions.

CAGE analysis can detect TSSs at single base-pair resolution, and therefore, it can be used to estimate the architecture of the promoter (Raborn et al., 2016). Shape index(SI) is one of the major indices used to evaluate promoter architecture (Hoskins et al.,2011). “Narrow” promoters initiate transcription at specific positions, while “broad”promoters initiate transcription at more dispersed regions. It is widely accepted that the promoter shape differs among different tissues or conditions (Forrest et al., 2014; Lizio et al., 2017). To detect promoter dynamics in the skeletal muscle under different metabolic conditions, we analyzed the promoter shape of each of the detected promoters in the reversible, hypometabolic, and torpor-specific groups (Figure 4E). In the torpor-specific groups, the down-regulated promoters showed a significantly different shape when compared to all muscle promoters (Figure 4F), while the GC richness did not show adifference (Figure S4E).

The torpor-specific promoters we found may represent regulators both upstream and downstream of the torpor transcriptional network. To further elucidate the early events involved in torpor-specific metabolism in peripheral tissues, it was necessary to place the animal in a condition where it had an unusually strong tendency to enter torpor, and to compare the muscle gene expression with that of normal torpor entry. For this, we mimicked the classical technique, sleep deprivation, which is frequently used in basic sleep research (Tatsuki et al., 2016; Wang et al., 2018), and performed torpor deprivation by gently touching the animal. Even when the mouse was not allowed to enter torpor, the *VO*_*2*_was close to that of Mid-torpor animals (Figure S4F). Furthermore, the transcriptome profile in the muscles from torpor-deprived animals did not show a clear difference from Mid-torporanimals in MDS plots (Figure S4G). When compared to Mid-torpor muscles, the torpor-deprived muscles had 45 up- and 27 down-regulated promoters (Figure 4G). Among these72 torpor-deprivation-specific promoters, one promoter starting at the minus strand ofchromosome 1: 191217941, namely the promoter of the activating transcription factor 3 (*atf3*) gene, was also found in the torpor-specific promoters (Figure 4H). Surprisingly, the binding site of ATF3 was one of the motifs enriched in the torpor-specific promoters (Figure S4H). The Atf3 motif was found in 33 of 289 torpor-specific promoters, and the peak of themotif probability was at 79 bp upstream of the TSS (Figure 4I).

These results showed that tissues of torpid mice have a torpor-specifictranscription signature. We also found that one of the torpor-specific genes, encodingtranscription factor ATF3 was more highly expressed during torpor deprivation. Furthermore, the ATF3-binding motif was found to be enriched in torpor-specific promoters. These findings were indicative of a novel pathway of active hypometabolism in peripheraltissues, possibly initiated by the torpor drive-correlated transcription factor ATF3. Finally, weanalyzed our promoter-based expression data with respect to the SNPs of B6J and B6N, tofind evidence that may explain the phenotypic difference between these two inbred strains.

### Genetic Link of Distinct Torpor Phenotypes in Inbred Mice

The classic laboratory mice B6J and B6N have very few genome differences, while theyshow distinct torpor phenotypes (Figure 1K). We discovered that the muscles in B6J miceshow torpor-specific expressions (Figure 4A). Because our data were analyzed by CAGE-seq, the promoter information, which is usually non-coding sequences, is directly available. Because most SNPs are found in non-coding regions, it is reasonable to analyze the SNPenrichment at the promoter regions of the torpor-specific expressions to explain theB6J/B6N difference.

First, we tested whether the 13 torpor-specific pathways (Figure 4D) were affectedby SNPs that are different between B6J and B6N (B6J/B6N). The SNPs located in thepromoter region of the genes included in the pathways were counted, and the enrichment was compared to the baseline to test the significance (Figure 5A). All 13 pathways showedsignificant enrichment (*p* < 0.05) indicating that the SNPs in B6J/B6N are strongly involvedin the torpor-specific pathways.

**Figure 5.**
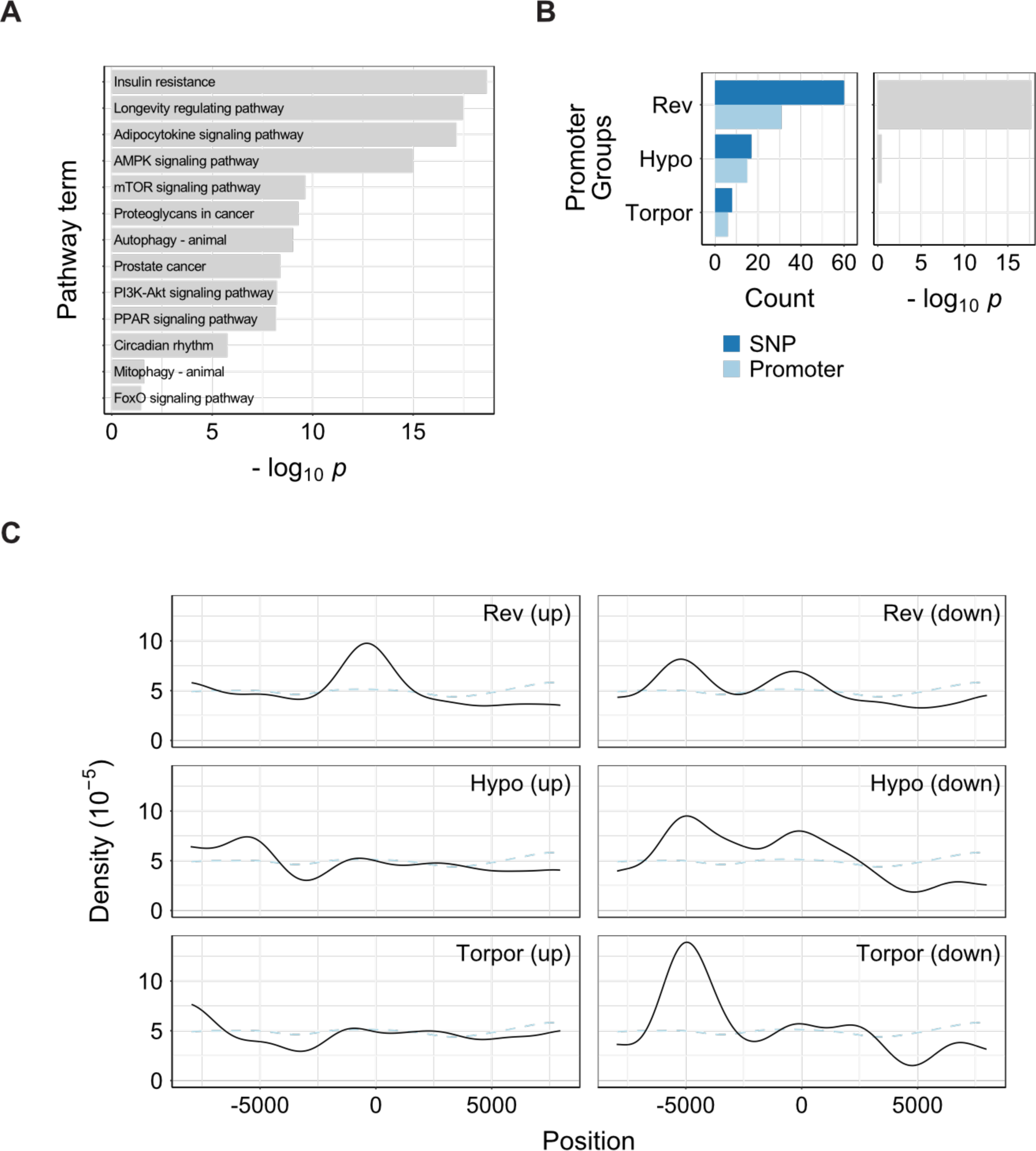
Genetic Link of Distinct Torpor Phenotypes in Inbred Mice. (A) Enrichment study of the B6J/B6N SNPs in torpor-specific promoter enriched KEGGpathways. (B) SNP counts in SNP-positive promoters (left). No group showed a significant enrichmentby B6J/B6N SNPs (right). Rev, Hypo, and Torpor denote reversible, hypometabolic, andtorpor-specific promoters. (C) SNP density was estimated in each promoter group. Light blue dashed line denotes thebackground SNP density.

Next, we tested the enrichment of SNPs at each promoter group. There were twoup- and four down-regulated promoters in the torpor-specific group that had at least oneSNP (Figure 5B). Possibly due to the low number of SNPs in this dataset, we were not ableto confirm a significant enrichment in torpor-specific promoters. The detailed position of theSNP in six promoters; *Plin5* and *Sik3* as up-regulated and *Creb3l1*, *Bhlhe40*, *Rrad*, and*Lrn1* as down-regulated promoters, are illustrated in Figures S5A and S5B.

Finally, we tested how the SNPs were distributed in the promoter region in eachgroup. We calculated the SNP density at a given position from the TSS (Figure 5C). Theresults indicated that SNPs tended to be enriched 5 kbp upstream from the TSS of torpor-specific down-regulated promoters.

These results collectively indicated that B6J/B6N SNPs may explain the torporphenotype difference in these two strains. In particular, the SNPs that were highly enrichedin the torpor-specific pathways designated the possible origin of the dissimilar torporphenotypes.

## DISCUSSION

### Mouse Torpor as a Model System for Active Hypometabolism

One goal of this study was to introduce mouse torpor as a study model for active hypometabolism. Hibernation is the most extreme phenotype of active hypometabolism,and there is a physiological distinction between hibernation and daily torpor (Ruf and Geiser, 2015). We recently showed that mouse torpor shares a common thermoregulation mechanism with hibernation in which the sensitivity of the thermoregulatory system is reduced (Sunagawa and Takahashi, 2016).

In this study, we extended our previous work by evaluating another inbred strainB6N. Despite the close genetic distance between B6N and B6J, we found that they had distinct torpor phenotypes (Figures 1F and S1E) due to a difference in heat production sensitivity (Figure S1F). Various inbred strains are reported to have distinct phenotypes,indicating a genetic involvement in torpor phenotypes (Dikic et al., 2008). Our findings strengthen this idea, because B6N and B6J have a very small genetic difference but a clear difference in torpor phenotypes. We also showed that the inbred strain-specific torpor phenotype is inheritable (Figures 1K and S1J), further validating the link between genetic background and torpor phenotype.

### Torpor-specific Transcriptions Differ from those of Hibernation and Starvation

In this study, we identified 287 torpor-specific promoters in mouse skeletal muscle (Figure 4A). Specificity was assured by including both reversible and hypometabolic promoters (Figures 2A and 3A). The results enabled us to identify likely metabolic pathways that are enriched during torpor (Figure 4D).

Circadian rhythm was the most enriched KEGG pathway by torpor-specific promoters (Figure 4D). The circadian clock is important in organizing metabolism and energy expenditure (Tahara and Shibata, 2013). In our study, the core circadian clock gene *per1* was up-regulated torpor-specifically, and *arntl1* was up-regulated in the reversible experiment. Because *per1* and *arnlt1* are normally expressed in reversed phases, our results in which both components were up-regulated together indicated that the circadian clock was disrupted in the skeletal muscle during torpor. Most past studies have focused onthe involvement of the central circadian clock (Ikeno et al., 2017; Revel et al., 2007), whilelittle is known about the peripheral circadian clock in torpid animals (Jansen et al., 2016). Thus, our results may provide evidence that the peripheral clock is disrupted during activehypometabolism.

Similarities between fasting during hibernation or daily torpor and calorie restrictionin non-hibernating mammals are reported (Xu et al., 2013b). During long-term torpor, suchas in hibernating mammals, carbohydrate-based metabolism switches to lipid use. Manystudies have suggested that the activation of AMPK is important in torpor induction (Lanaspa et al., 2015; Melvin and Andrews, 2009; Zhang et al., 2015). However, anotherstudy demonstrated AMPK activation only in white adipose tissue, not in the liver, skeletalmuscle, brown adipose tissue, or brain, during hibernation (Horman et al., 2005). Our studycorroborates the findings of Horman’s research, by demonstrating no significant changes inthe AMPK-encoding gene expression during torpor in skeletal muscle.

The PPAR-signaling pathway also regulates lipid metabolism. Numerous studieshave shown increased PPARs in various organs at the mRNA and protein levels duringtorpor, in several hibernating species (Han et al., 2015; Xu et al., 2013b). Recently, an over-expression of PPARα protein in mouse liver, comparable to that in hibernating bats wasreported, suggesting a potential hibernation capability of mice (Han et al., 2015). Accordingto our data, *Pparα* is upregulated in torpid mice muscle along with several target genesassociated with cholesterol metabolism and fatty acid transport. Remarkably, *Ppargc1a* gene, encoding PGC-1α (peroxisome proliferator-activated receptor-γ coactivator-1) wasalso over-expressed in mice during torpor. Recently, PGC-1α activation was suggested tobe responsible for protecting skeletal muscle from atrophy during long periods of torpor inhibernators (Xu et al., 2013a). Our results suggest that a similar pathway may be activatedin mouse torpor as well.

We found that the insulin/Akt and mTOR signaling pathways, which have roles inskeletal muscle remodeling and metabolic rate depression, were enriched. Previous studiesshowed that insulin signaling is inhibited in the skeletal muscle of torpid gray mouse lemurs (Tessier et al., 2015) and that the Akt kinase activity is suppressed during torpor in multiple tissues of ground squirrels (Abnous and Storey, 2008; Cai et al., 2004; Wu and Storey,2012). The suppressed Akt activity is accompanied by a reduction in mTOR activation,leading to a state of protein synthesis inhibition during torpor in hibernators (Lee et al.,2010; McMullen and Hallenbeck, 2010; Wu and Storey, 2012). Our results demonstrated adown-regulation of *igf1*, which encodes IGF-1, and an activation of *mtor*, which encodes mTOR, in torpor, which appear to be paradoxical to past studies.

The Insulin/Akt pathway also controls the phosphorylation and activation of theFOXO1 transcription factor, a disuse atrophy signature that upregulates the muscle-specific ubiquitin ligases *trim63* (MuRF1) and *fbxo30* (Atrogin-1). In our study, we found that FOXO1, MuRF1, and Atrogin-1 were up-regulated, as in the case of disuse atrophy in mice and rats (Sandri et al., 2004; Senf et al., 2010).

In summary, we found that the up-regulation of PGC-1α and down-regulation ofIGF-1 in the skeletal muscle of torpid mice are similar to hibernating animals, in which they contribute to muscle protection and the suppression of protein synthesis. On the otherhand, muscle atrophy and autophagy signatures such as FOXO1, MuRF1, and Atrogin-1were up-regulated during torpor, indicating that atrophic changes is also progressed. Furthermore, mTOR activation was found, which is a signature of muscle hypertrophy. Thus, we can conclude that mouse torpor has a unique transcription profile, sharing signatures with hibernation, starvation-induced atrophy, and muscle hypertrophy.

### Dynamics of Torpor-specific Transcriptions

The deep CAGE technology enabled us to evaluate the dynamics of the torpor-specific promoters. We found that down-regulated torpor-specific promoters were narrower than other muscle promoters (Figures 4E and 4F). However, the GC content of the torpor promoters was not significantly different from that of all muscle promoter regions (Figure S4E): approximately half of the TSSs were located in CpG islands, in which both AT- and GC-rich motifs were overrepresented (Figures S4C and S4D).

To gain insight about the upstream network of torpor, we evaluated a torpor-deprived condition. Note that this dynamic state, the torpor-deprived condition, is verydifficult to induce in hibernators, because very little stimulation can cause them to halttorpor induction. Taking advantage of this torpor-deprivation state in mice, we identifiedtranscription factor ATF3 as a candidate factor that is correlated with the need to entertorpor. *Atf3* is a well-known stress-inducible gene (Hai and Hartman, 2001). Recentcumulative evidence suggests ischemia/reperfusion significantly induce ATF3 expression invarious organs (Lee et al., 2013; Rao et al., 2015; Yoshida et al., 2008). In the currentstudy, *Atf3* was identified not only as a stress-induced gene, but also as a torpor-drivecorrelated factor. Torpor is an active-hypometabolic condition, which can be described as aphysiological ischemia. Although we lack direct evidence, we propose the hypothesis that *Atf3* may be a factor mediating the initiation of hypometabolism, and because of that, it isexpressed to protect the organs under stressful conditions such as ischemia.

Another advantage of deep CAGE is the rich information obtained about thepromoter region of the expression of interest. This study exploited our finding that B6J andB6N have different torpor phenotypes. To identify which SNP was responsible for thephenotype difference, we used all of the data acquired in this study. Although the down-regulated torpor-specific promoters tended to have more SNPs, we were unable to identifyspecific SNPs related to the torpor phenotype from observation (Figure 5B). Therefore,further study is needed to test how the candidate SNPs (Figures S5A and S5B) affect thetorpor phenotype by genetic intervention.

### Fundamental Understanding of Active Hypometabolism for Medical Applications

The overall results of this study indicate that the mouse is an excellent animal for studyingthe as-yet-unknown mechanisms of active hypometabolism. Understanding the core engineof the hypometabolism in torpid tissues will be the key to enabling non-hibernating animals,including humans, to hibernate. Inducing active hypometabolism in humans would be animportant breakthrough for many medical applications (Bouma et al., 2012). The benefits tousing mice are not limited to technological advances in genetics, but extend to theenormous potential for *in vitro* studies using cell or tissue culture. In stem cell biology, patient-derived stem cells represent a valuable resource for understanding diseases anddeveloping treatments, because the cells reflect the phenotype of the patient (Avior et al.,2016). We believe, similarly, that mouse-derived stem cells or tissues will provide a uniqueplatform for investigating strain-specific hypometabolic phenotypes in animals. Moreover,because *in vitro* studies can be easily extended to experiments using human cells/tissuederived from human induced pluripotent stem cells, active hypometabolism research inmouse cells/tissues is an important step toward the realization of human hypometabolism.

## EXPERIMENTAL PROCEDURES

### Animals

All animal experiments were performed according to the guidelines for animal experiments of the RIKEN Center for Biosystems Dynamics Research and approved by the Animal Experiment Committee of the RIKEN Kobe Institute. C57BL/6NJcl mice were purchased from CLEA Japan, Inc. and C57BL/6J mice were from Oriental Yeast Co., Ltd. Until the mice were used in torpor experiments, they were given food and water ad libitum and maintained at a *T*_*A*_ of 21 °C, a relative humidity of 50%, and with a 12-hr light/12-hr darkcycle. The *T*_*B*_ and the *VO*_*2*_ of the animal were continuously recorded by an implanted telemetry temperature sensor (TA11TA-F10, DSI) and by respiratory gas analysis (ARCO-2000 mass spectrometer, ARCO system), respectively. See SUPPLEMENTALEXPERIMENTAL PROCEDURES for details.

### Non-Amplified non-Tagging Illumina Cap Analysis of Gene Expression (nAnT-iCAGE)Library Preparation and Sequencing

RNA was isolated from sampled tissues (see SUPPLEMENTAL EXPERIMENTALPROCEDURES). Transcriptomics libraries were prepared according to a standard protocol for the CAGE method by using 5 μg of extracted total RNA from mouse muscles (Murata etal., 2014). The RNA was used as a template for the first strand cDNA synthesis, which was then biotinylated at the 5’-end to allow streptavidin capture. Linkers were then attached at the 5′ and 3′ ends, and the second strand cDNA was synthesized. The quality of the libraries was verified using a Bioanalyzer 2100 (Agilent), and the yield was validated by qPCR. The single-end libraries were then sequenced on a NextSeq platform (Illumina) or on a HiSeq 250 platform using Rapid Run mode (Illumina), in experiment #1 and #2,respectively.

### Mapping, Peaks Calling, and Annotation

Sequenced reads were trimmed and mapped on the mouse mm10 genome assembly using bwa and hisat2 (Kim et al., 2015; Li and Durbin, 2010). For each sample, we obtained CAGE-defined TSSs (CTSSs) according to the reads abundance, and then clustered themusing PromoterPipeline (Arnaud et al., 2016), the highest peaks were annotated as TSSs. These CAGE clusters were then associated with their closest genes using the Ensembl andRefseq transcripts annotation available for mm10. The accession number for thesequencing data reported in this work is GEO: GSE117937.

## ACKNOWLEDGMENTS

We thank the LARGE, RIKEN BDR for housing the mice. This work was supported by theRIKEN Special Postdoctoral Researcher program (G.A.S.) and by Grant-in-Aid for ScientificResearch on Innovative Areas (Thermal Biology) 18H04706 from MEXT (G.A.S.).

## AUTHOR CONTRIBUTIONS

G.A.S., O.G., and M.T. designed the study. G.A.S and K.I. performed the animalexperiments and tissue sampling, supervised by G.G. R.D. and G.A.S. analyzed the data.G.A.S., R.D., G.G., and G.O. wrote the manuscript. All authors discussed the results andcommented on the manuscript text.

## ADDITIONAL INFORMATION

### Competing financial interests

The authors declare no competing financial interests.

## SUPPLEMENTAL EXPERIMENTAL PROCEDURES

### Animals

For the C57BL/6NJcl mice, 58 total mice were used to characterize their torpor phenotype(12 females and 46 males). The age (mean ± SD) and body weight (mean ± SD) at the beginning of the experiment were as follows: 8.67 ± 0.39 weeks old and 19.6 ± 0.7 g for females; 8.34 ± 0.53 weeks old and 22.5 ± 1.2 g for males. For the C57BL/6J mice, 50 total mice were used to characterize the torpor phenotype (n = 12; 8 females and 4 males) and sampling tissues (n = 38, all males). The characteristics were: 8.43 ± 0.15 weeks old and17.7 ± 0.6 g for females; 8.22 ± 0.39 weeks old and 23.0 ± 1.2 g for males. As described inthe RESULTS section, for B6J mice, data recorded in a previous report (Sunagawa and Takahashi, 2016) (n = 43, all male mice, 8.07 ± 0.35 weeks old, 22.9 ± 1.2 g) were also included in the data analysis to characterize the thermoregulatory system. To test for torpor phenotype inheritance, B6J and B6N were crossed, and their offspring were evaluated. Male B6NJ mice (B6N females crossed with B6J males) and B6JN mice (B6J females crossed with B6N males) were used for this assessment. The characteristics of each strain 16 were: n = 9, 8.90 ± 1.01 weeks old, 25.5 ± 2.9 g for B6NJ-F1 mice and n = 8, 8.73 ± 0.62weeks old and 23.3 ± 0.9 g for B6JN-F1 mice.

During the experiments, each animal was housed in a temperature-controlled chamber (HC-100, Shin Factory). To record *T*_*B*_ continuously, a telemetry temperature sensor (TA11TA-F10, DSI) was implanted in the animal’s abdominal cavity under general inhalation anesthesia at least 7 days before recording. The metabolism of the animal was continuously analyzed by respiratory gas analysis (ARCO-2000 mass spectrometer, ARCO system). During the experiment, the animal was monitored through a networked videocamera (TS-WPTCAM, I-O DATA, Inc.). This video camera can detect infrared signals,which made it possible to monitor the animal’s health during the dark phase without opening the chamber.

### Daily Torpor Induction Experiment

Each daily torpor induction experiment was designed to record the animal’s metabolism forthree days (Figure 1B) unless the tissues were sampled on day 2. The animals were introduced to the chamber the day before recording started (day 0). Food and water were freely accessible. The *T*_*A*_ was set as indicated on day 0 and kept constant throughout the experiment. A telemetry temperature sensor implanted in the mouse was turned on before placing the mouse in the chamber. The standard experimental design was as follows: on day 2, ZT-0, the food was removed to induce torpor. After 24 hours, on day 3, ZT-0, the food was returned to each animal. In the torpor-prevention experiment with food administration (Figure 3A), the food was not removed at day 2. In the torpor-deprivation experiment, one experimenter monitored the *VO*_*2*_ and touched the mouse gently when the *VO*_*2*_ started todrop. The metabolism monitoring for torpor deprivation was started at ZT-17 on day 2 and maintained until the mouse tissue was sampled at ZT-22.

### Body Temperature and Oxygen Consumption Modelling for Daily Torpor Detection

To model the temporal variation of *T*_*B*_ and *VO*_*2*_, we constructed the models in a Bayesian framework. From the first 24-hour recordings of *T*_*B*_ and *VO*_*2*_ for each animal, we estimated the parameters using Markov Chain Monte Carlo (MCMC) sampling by Stan (StanDevelopment Team, 2016a) with the RStan library (Stan Development Team, 2016b) in R (R Core Team, 2017). The detailed methods were described previously (Sunagawa andTakahashi, 2016) and modified with software updates. In short, we used the 99.9% credible interval (CI) of the posterior distribution of the estimated metabolism, the *T*_*B*_ and *VO*_*2*_, of theanimal to detect outliers. That is, when the value was lower than the CI, that time point was defined as torpor due to an abnormally low metabolic status. In this study, when both *T*_*B*_and *VO*_*2*_ met the criteria in the second half of the day, the time point was labelled as torpor.

### Parameter Estimation of the Thermoregulatory system

The thermoregulatory system was modelled as an integration of the heat loss and heat production of the animal. Three parameters *G*, *T*_*R*_, and *H* were estimated from the metabolic stable state of the animal at various *T*_*A*_s. The details were described previously (Sunagawa and Takahashi, 2016).

### Tissue Sampling and RNA Isolation

Dissected soleus muscles were rapidly frozen in liquid nitrogen. The RNA was isolated using an RNeasy Fibrous tissue kit (Qiagen) according to the manufacturer’s instructions. The quality of the total RNA was evaluated using a Bioanalyzer 2100 (Agilent). The quantity and purity of the RNA were estimated using a NanoDrop Spectrophotometer. The lateral or both soleus muscles were used according to the total amount of RNA needed.

### Data Processing

Data were processed in R (R Core Team, 2017) unless otherwise noted. The expressionlevel of the 12,862 defined CAGE clusters was normalized by sample in TPM (tags permillion) and then analyzed with the edgeR package (Robinson et al., 2010) with TMM(trimmed mean of M-value) normalization. For MDS (multidimensional scaling) plots, DE(differential expression), and GO and KEGG pathway enrichment analysis, several Rpackages were applied, including edgeR, clusterProfiler (Yu et al., 2012), and pathview (Luo and Brouwer, 2013). Muscle enhancers were predicted de novo by applying theFANTOM5 protocol (Andersson et al., 2014) to our mouse CAGE data and masked with±500-bp regions from the 5’ ends of annotated genes. The mouse CAGE data for muscles can be observed and are publicly available in the Zenbu browser(http://fantom.gsc.riken.jp/zenbu/gLyphs/#config=ylDd70XVLdPufetrnXzQkB). The DEresults (reversible, hypometabolic, and torpor-deprivation-specific promoters) along with torpor-specific promoters are listed in Supplemental Table S1.

### Basic Promoter Features Analysis

Promoter region features were analyzed in terms of GC content and SI (Hoskins et al.,2011). The SI and %GC were calculated for ±50 bp regions around the TSS position. CpGisland muscle promoters were defined by searching for overlaps with the UCSC annotation using bedtools v2.25.

### Motif Analysis

Transcription factor binding sites (TFBS) were predicted in −300/+100 bp regions around theTSS position using MEME Suite 4.11.2 and the JASPAR CORE motif library for vertebrates2016. The position-dependent enrichment of these motifs was performed by the CentriMotool.

### SNP analysis

The single nucleotide polymorphisms data for the C57BL/6NJ strain were downloaded fromthe Mouse Genomes Project of the Sanger Institute (ftp://ftp-mouse.sanger.ac.uk/current_snps/strain_specific_vcfs/C57BL_6NJ.mgp.v5.snps.dbSNP142.vcf.gz). Originated from the C57BL/6N strain, the C57BL/6NJ mice were derived fromembryos cryopreserved (F126) at the NIH in 1984, and C57BL/6NJcl mice were introducedto the Central Institute for Experimental Animals from the NIH at F121 in 1978, and thentransferred to CLEA Japan at F146 in 1988. Of the C57BL/6NJ-specific SNPs, 89% arepreserved in C57BL/6NJcl (Mekada et al., 2015). Overlaps of the SNPs with mousetranscripts, muscle promoters, and enhancers regions were performed using bedtoolsv2.25. All SNPs overlapping predicted promoter regions (−300/+100 bp from TSS),annotated RefSeq and Ensembl transcripts, including both coding and noncoding regions,were counted. The overrepresentation rate of SNPs in pathways was calculated by applyinga hypergeometric test in R.

## SUPPLEMANTAL DATA

**Figure S1.**
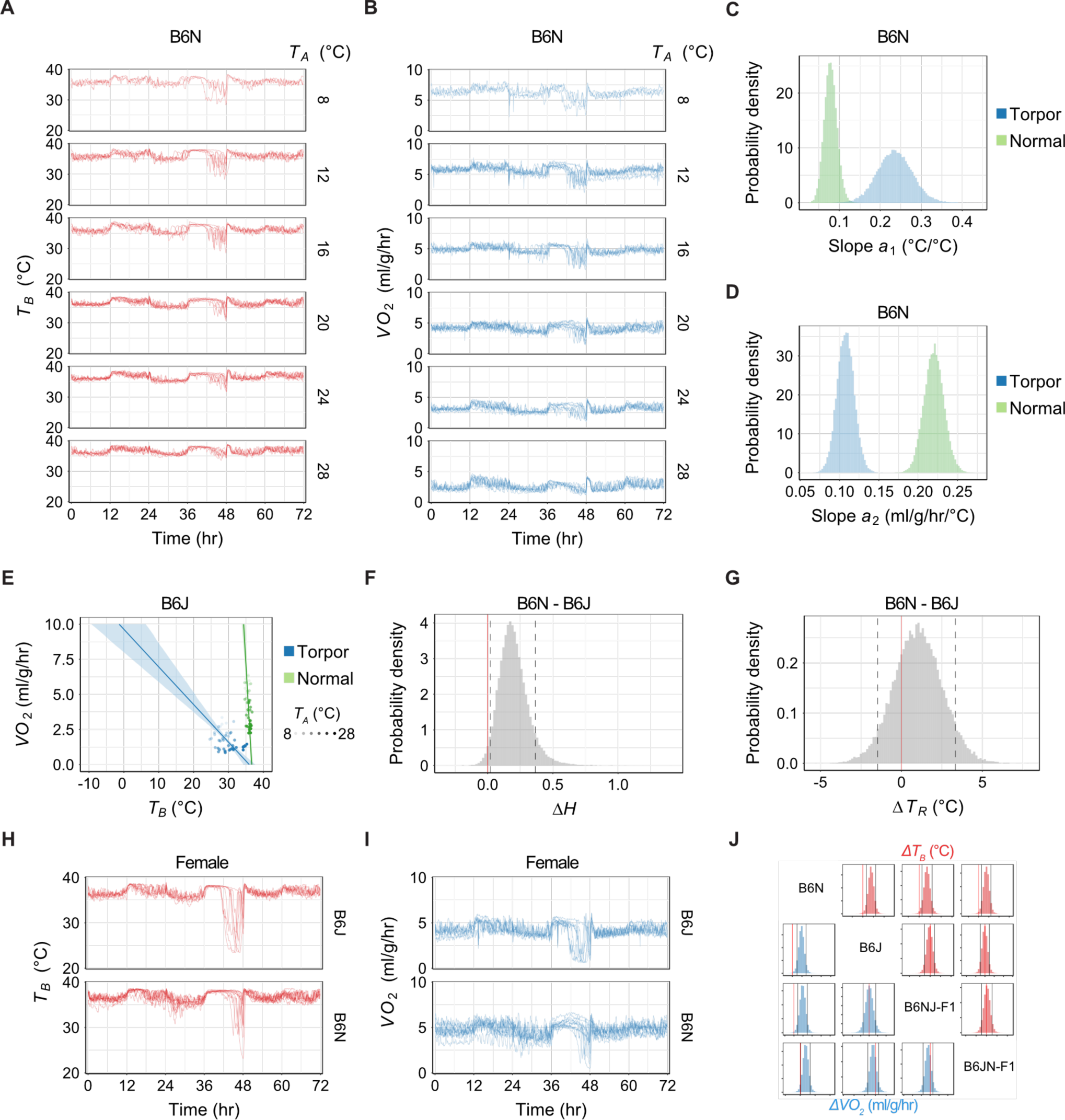
Torpor Phenotype in Mice is Affected by Genetic Background, related to Figure 1. (A) (B) Traces of *T*_*B*_ (red lines) and *VO*_*2*_ (blues lines) of the B6N male mouse at various *T*_*A*_s. (C) Posterior distribution of the estimated slope *a*_*1*_ during the normal state and torpor. (D) Posterior distribution of the estimated slope *a*_*2*_ during the normal state and torpor. (E) Relationship between minimum *T*_*B*_ and *VO*_*2*_ seen during the normal and torpid states atvarious *T*_*A*_s in B6J mice. Darkness of the dots indicates the *T*_*A*_. The horizontal intercept of the line indicates the theoretical set-point of *T*_*B*_, which is *T*_*R*_. Note that the slope of the *T*_*B*_ -*VO*_*2*_ relationship during torpor was less steep for B6J than for B6N mice, indicating that B6Jhad less sensitivity to *T*_*B*_ during torpor, consistent with the observation that B6J had a lowerminimal *T*_*B*_ during torpor than B6N. (F) Distribution of the estimated *ΔH*, the difference in *H* during torpor for B6N versus B6J. Red line denotes 0, and the dashed lines denote the lower and upper range of the 89%HDPI of *ΔH*. Note that because the HDPI does not include 0, the *ΔH* is likely to be positive at the probability of more than 89%; this can be interpreted as it being highly probable thatB6N has a larger *H* than B6J. (G) Distribution of the estimated *ΔT*_*R*_, the difference in of *T*_*R*_ during torpor of B6N versusB6J. Red line denotes 0, and dashed lines denote the lower and upper range of 89% HDPIof *ΔT*_*R*_. Because the 89% HDPI do includes 0, the *ΔT*_*R*_ may be 0 at a probability of 89%;this can interpreted as it being highly probable that B6N and B6J do not have different *T*_*R*_s. (H) (I) Traces of *T*_*B*_ (red lines) and *VO*_*2*_ (blues lines) over time of B6J and B6N female miceat *T*_*A*_ = 20 °C. (J) Posterior distribution of the difference in the estimated minimal *T*_*B*_ and *VO*_*2*_ of B6N, B6J,B6NJ-F1, and B6JN-F1 mice during torpor. Red vertical line denotes 0, and the dashed vertical lines denote the lower and upper range of the 89% HDPI of *ΔT*_*B*_ or *ΔVO*_*2*_. When 0 isnot included in the HDPI, the index is highly probable to have a difference. B6N had adistinct phenotype for both the minimal *T*_*B*_ and *VO*_*2*_ during torpor than from that of the other three strains.

**Figure S2.**
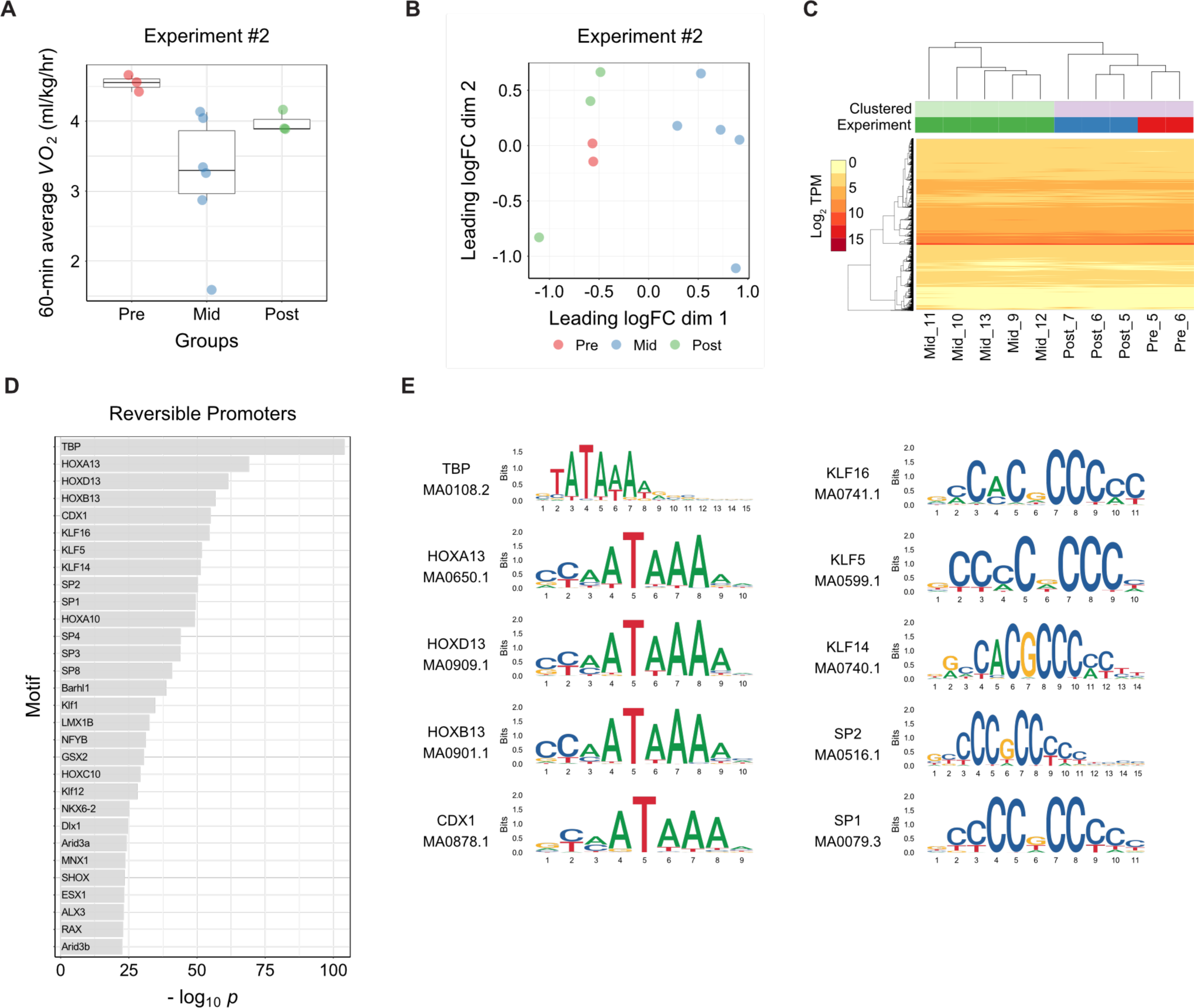
Fasting-induced Torpor Shows a Reversible Transcriptome Signature,related to Figure 2. (A) Boxplots for the *VO*_*2*_ of animals at sampling in reversibility experiment #2. Each dot represents one sample from one animal. The results resembled the metabolic phenotypes as detected in experiment #1. See Figure 2B. (B) MDS plot of the TSS-based distance in reversibility experiment #2. Each dot represents one sample from one animal. Note that the Mid group was clustered differently from the Preand Post groups in the 1st dimension, as it were in Figure 2C. (C) Hierarchical clustering heatmap based on the TPM of TSS detected in the reversibility experiment #2. (D) The top thirty motifs enriched in the reversible promoters. (E) Logos of the top ten motifs in the reversible promoters.

**Figure S3.**
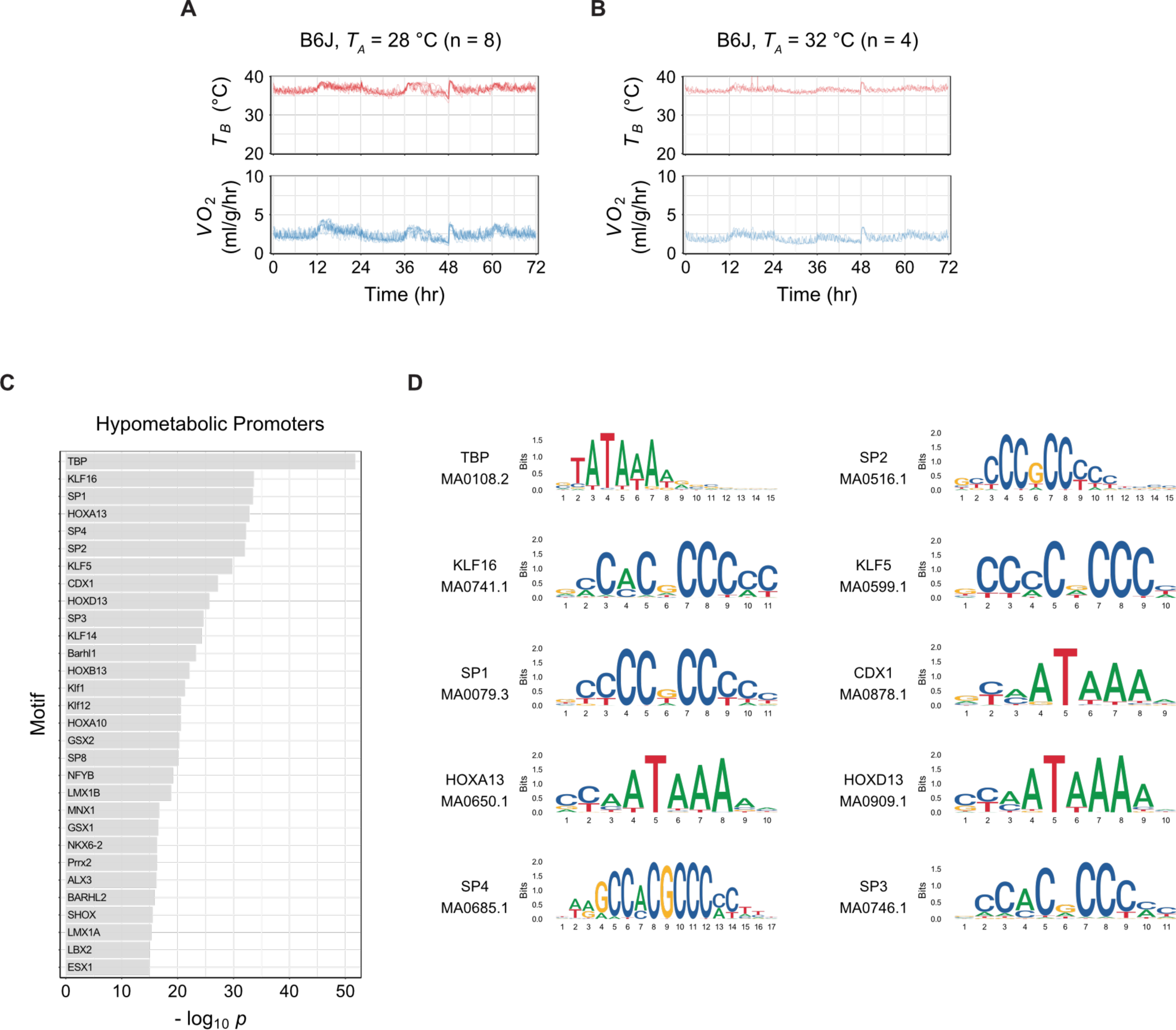
Torpor Prevention at High *T*_*A*_ Revealed Hypometabolism-associatedPromoters, related to Figure 3. (A) One B6J mouse in eight failed to enter torpor at *T*_*A*_ = 28 °C. (B) At *T*_*A*_ = 32 °C, no mouse entered torpor (n = 4). (C) Top thirty motifs enriched in the hypometabolic promoters. (D) Logos of the top ten motifs enriched in the hypometabolic promoters.

**Figure S4.**
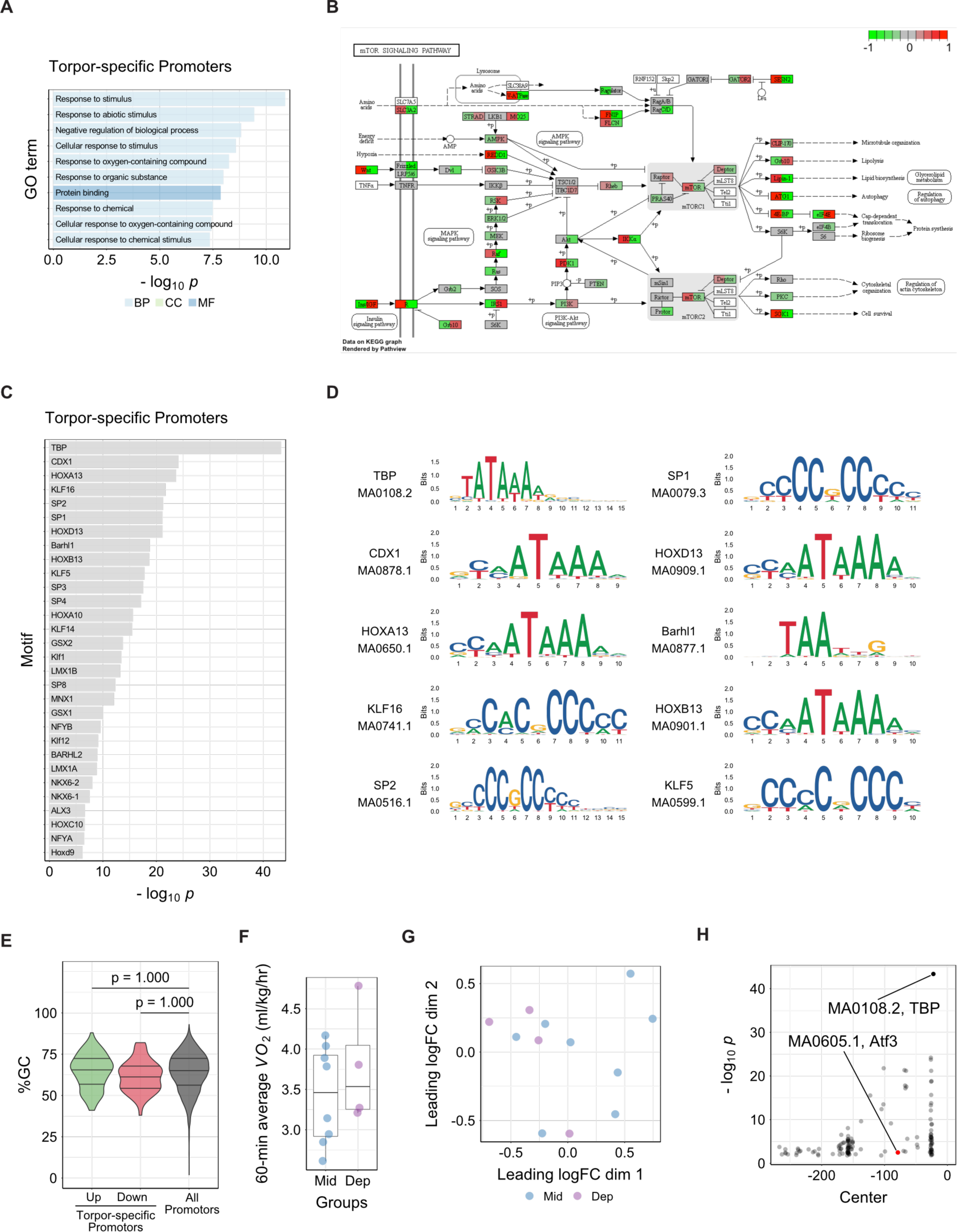
Identification of Torpor-specific Promoters and their Dynamics, related to Figure 4. (A) Top ten enriched GO terms in the torpor specific promoters. (B) Of the 13 enriched KEGG pathways, the “mTOR signaling pathway” is shown as arepresentative example. Green and red denote up- and down-regulated genes,respectively. (C) Top thirty motifs enriched in the torpor-specific promoters. (D) Logos of the top ten enriched motifs in the torpor-specific promoters. (E) Distribution of the %GC in the torpor-specific promoters compared to all muscle promoters. The three horizontal lines inside the violin denote the 1st, 2nd, and 3rd quartile of the distribution from the upmost line. No significant difference was detected in this dataset. (F) Boxplots for the *VO*_*2*_ of animals at sampling in the torpor deprivation experiment. Each dot represents one sample from one animal. Torpor-deprived animals (Dep group, n = 4)did not show an apparent change in *VO*_*2*_ compared to the Mid group. (G) MDS plot of the TSS-based distance in the torpor-deprivation experiment. Each dot represents one sample from one animal. A clear separation between the Mid and Dep groups was not found in this analysis. (H) Distribution of motifs enriched in the torpor-specific promoters. The horizontal axis denotes the position of the motif density peak from the TSS. The vertical axis denotes the*p*-value of the enriched motif.

**Figure S5.**
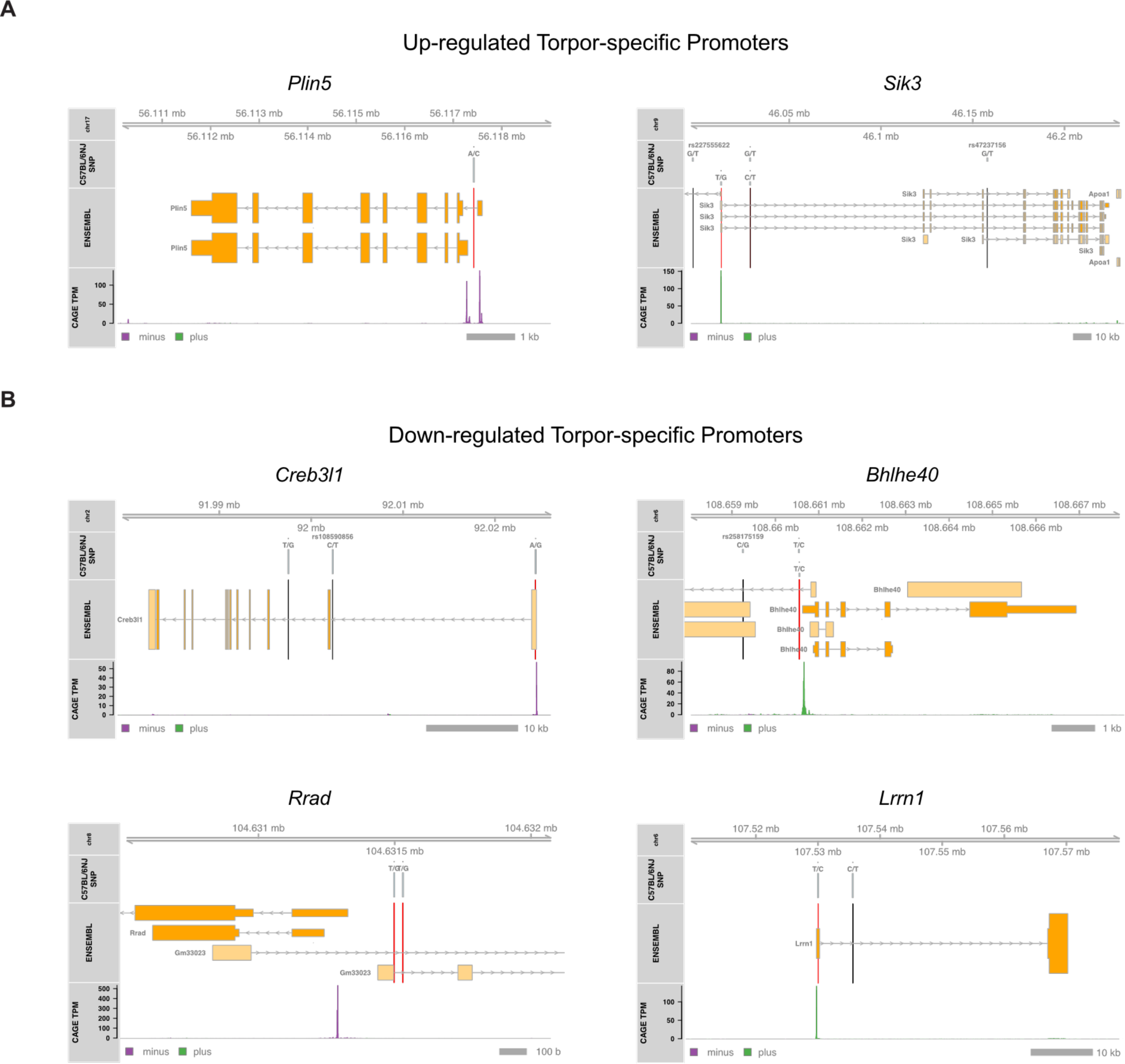
Genetic Link of Distinct Torpor Phenotypes in Inbred Mice, related toFigure 5. (A) (B) Torpor-specific promoters that have B6N/B6J SNPs within the range of +400/-100bp from the TSS. The vertical lines denote the SNPs, which are red when included in thepromoters and black when not. Among the up-regulated torpor-specific promoters, *Plin5*and *Sik3* had one SNP each. Among the down-regulated promoters, *Creb3l1* and *Lrrn1* hadone, and *Bhlhe40* and *Rrad* had two SNPs in the promoter region.

**Table S1.** Differentially Expressed Promoters During Torpor

